# Implications of successive blood feeding on *Wolbachia*-mediated dengue virus inhibition in *Aedes aegypti* mosquitoes

**DOI:** 10.1101/2025.02.06.636928

**Authors:** Rebecca M. Johnson, Mallery I. Breban, Braiya L. Nolan, Afeez Sodeinde, Isabel M. Ott, Perran A. Ross, Xinyue Gu, Nathan D. Grubaugh, T. Alex Perkins, Doug E. Brackney, Chantal B. F. Vogels

## Abstract

Dengue virus (DENV) is a mosquito-borne virus that poses a continued and increasing threat to public health. A promising strategy to mitigate the burden of DENV is introgression of the virus-inhibiting *Wolbachia pipientis* bacterium into *Aedes aegypti* populations in the field. While previous studies on *Wolbachia*-mediated virus inhibition have typically assessed viral replication following a single bloodmeal, the main vector of DENV, *Ae. aegypti*, feeds frequently, often biting multiple hosts per gonotrophic cycle and promptly attempting to refeed following egg laying. Previously, we demonstrated that successive blood feeding reduces the extrinsic incubation period (EIP) and shortens the time it takes for a mosquito to be able to transmit viruses to a new host. With this in mind, we investigated the impact of successive blood meals on DENV serotype 2 (DENV-2) in *Ae. aegypti* in the presence or absence of *Wolbachia* (*w*AlbB and *w*MelM). We found that both WT and *Wolbachia* transinfected had increased DENV-2 dissemination 7 days post-infection as well as higher body titers of DENV-2 in the double-fed groups. Using these empirical data in a binomial regression model, we estimated that successive feeding increased the probability of WT and *Wolbachia* transinfected mosquitoes surviving the EIP. When we estimated the odds of surviving the EIP for mosquitoes with *Wolbachia* relative to WT mosquitoes, successive feeding increased the chances of WT mosquitoes surviving the EIP more than in mosquitoes with *Wolbachia*, indicating a strong inhibitory effect of *Wolbachia* even in the context of natural frequent blood feeding behavior. Our work shows that mosquito feeding behavior should be considered when assessing the inhibitory effects of *Wolbachia* on DENV.

## Main

Dengue virus (DENV) is a mosquito-borne virus that poses a significant public health threat, which is exemplified by the current outbreak and a record number of reported cases during the first half of 2024 alone^1,2^. To mitigate this increasing burden, novel control strategies are needed to disrupt DENV transmission between *Ae. aegypti* mosquitoes and humans. One such control strategy is the release of *Ae. aegypti* mosquitoes transinfected with different strains of the virus-inhibiting *Wolbachia* bacterium^3,4^. *Ae. aegypti* populations transinfected with different variants of the *Wolbachia w*Mel or *w*AlbB strains are currently released in the field to either suppress or replace local mosquito populations^5–8^. Mosquito population replacement strategies rely on the ability of *Wolbachia* to inhibit mosquito-borne virus transmission by mosquitoes, including DENV. However, virus inhibition can be incomplete, and the underlying mechanisms are not fully understood, which warrants further investigation into the underlying factors that may influence the efficiency of virus inhibition^9^.

Recently, we found that natural mosquito feeding behavior can influence virus dissemination (exit from the mosquito midgut to secondary tissues such as the salivary glands) and the extrinsic incubation period (EIP; the duration between virus acquisition and transmission)^10–13^. If mosquitoes feed frequently, as is often seen in *Ae. aegypti* in the wild, virus disseminates from the mosquito midgut faster resulting in a longer time period in which mosquitoes can transmit virus to susceptible hosts, and a larger projected R_0_ ^10,11,14–16^. Although some strains of *Wolbachia* can be highly effective in disrupting the transmission of DENV and other mosquito-borne viruses, many of these experiments did not offer mosquitoes more than one blood meal before assessing transmission ability and current modelling of *Wolbachia* efficacy frequently relies on dissemination times calculated from mosquitoes fed a single blood meal^4,17,18^. While previous studies with *Wolbachia*-infected mosquitoes have either provided successive noninfectious blood meals followed by an infectious feed, or provided successive infectious blood meals following an extended egg quiescence, no studies have determined the impact of successive feeding after an initial infectious blood meal^19,20^. In this study, we investigated the effect of successive blood feeding on the inhibition of DENV-2 in *Ae. aegypti* mosquitoes stably transinfected with *w*MelM and *w*AlbB *Wolbachia* strains. Specifically, we evaluated the hypothesis that successive blood feeding decreases the effectiveness of *Wolbachia* by facilitating more efficient DENV dissemination. Our work has implications for understanding *Wolbachia*-mediated virus inhibition, the use of *Wolbachia* transinfected mosquitoes for population replacement, and further DENV control efforts.

## Results

To test our hypothesis that successive feeding results in more efficient virus dissemination, we provided a DENV-2 spiked infectious blood meal to wild-type *Ae. aegypti* mosquitos lacking *Wolbachia* (WT), and *Aedes aegypti* mosquitoes stably transinfected with *w*MelM (*w*MelM) or *w*AlbB (*w*AlbB) followed by a second non-infectious blood meal in the “double-feed” group (**Fig 1a**). First, we compared infection and dissemination rates across colonies and feeding groups to establish *Wolbachia* inhibition phenotypes and the impact of successive blood meals. In both single-fed and double-fed groups, we found intermediate DENV-2 inhibition in *w*AlbB mosquitoes and stronger inhibition in *w*MelM mosquitoes as compared to those without *Wolbachia* (**Fig 1b-c**, p < 0.01). This is consistent with dengue inhibition phenotypes previously reported for these *Wolbachia* strains^3,21^. For WT mosquitoes, we observed no difference in infection rates between the single- and double-fed groups, although dissemination was increased in the double-fed group, consistent with our previous work (**Fig 1c**)^10^. For *w*AlbB, we found a similar effect with a significant increase in dissemination after the second blood meal (**Fig 1c**). While we found similar trends for *w*MelM, strong inhibition resulted in low numbers of infected mosquitoes and even lower numbers with subsequent dissemination and therefore we were unable to detect any significant differences between the *w*MelM single- and double-fed groups. Our findings show that successive blood feeding results in increased dissemination at 7 days post-infection (dpi) in both the presence and absence of *Wolbachia*.

**Figure 1:**
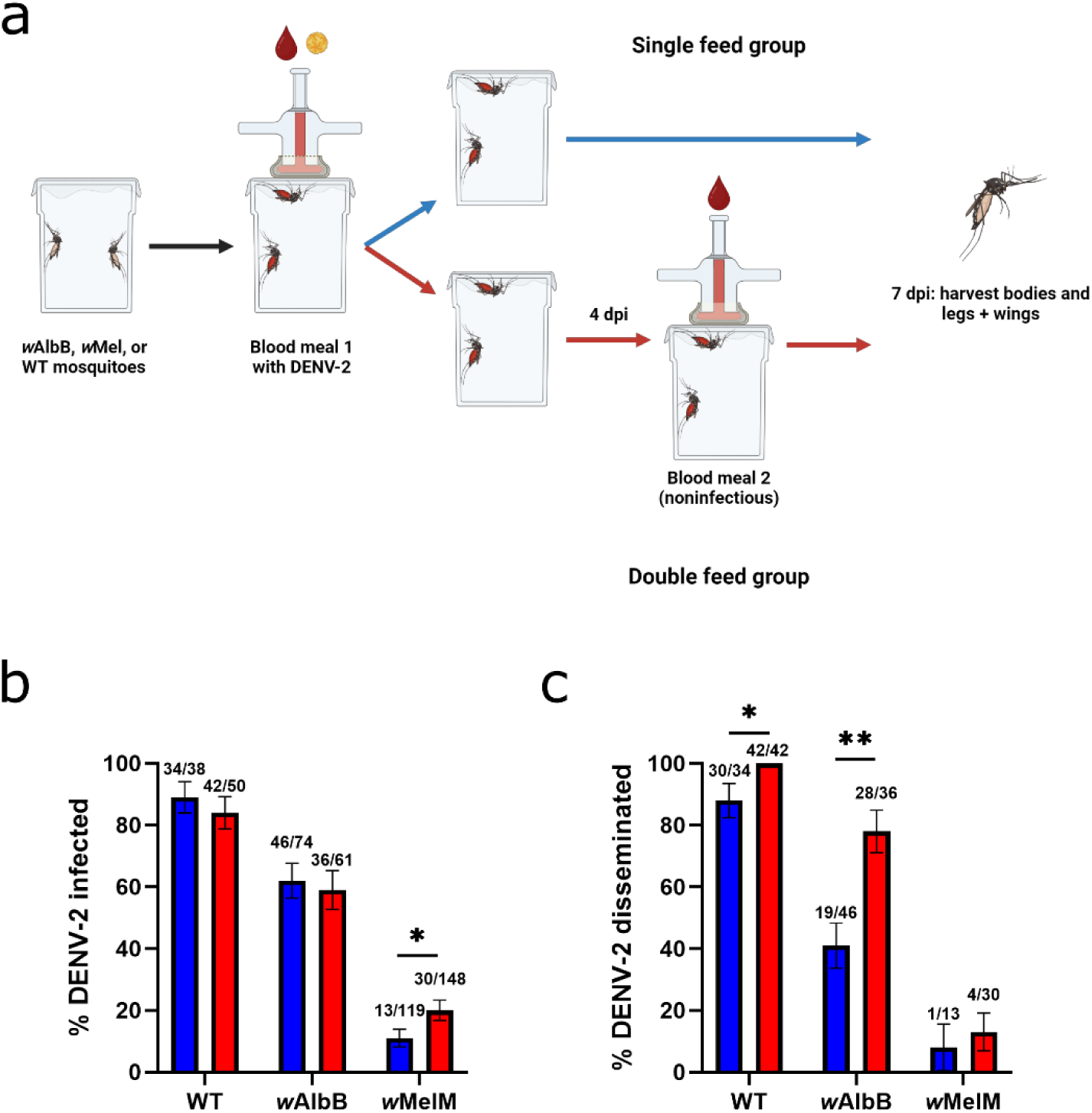
Transinfection with *w*AlbB or *w*MelM *Wolbachia* reduces DENV-2 infection in *Ae. aegypti* and successive feeding leads to higher rates of dissemination at 7 dpi. **a)** Experimental design for initial infection and dissemination studies of single- and double-fed WT, *w*AlbB, and *w*MelM mosquitoes. **b)** Proportion of infected single- and double-fed WT, *w*AlbB, and *w*MelM mosquitoes 7 dpi. **c)** Proportion of single- and double-fed WT, *w*AlbB, and *w*MelM mosquitoes with disseminated infection 7 dpi. Comparisons were made using Fisher’s exact tests. * = p ≤ 0.05, ** = p ≤ 0.01, *** = p ≤ 0.001, **** = p < 0.0001. Blue = single-fed, red = double-fed. Lines indicate mean **±** standard error of the mean.

To explore the basis of increased dissemination after successive blood feeding, we measured DENV-2 titers and *Wolbachia* densities across treatment groups. For WT and *w*AlbB groups, taking a second, noninfectious blood meal led to higher DENV-2 genome equivalents/μl as measured via qPCR, whereas we observed no significant differences in DENV-2 levels between single- and double-fed mosquitoes transinfected with *w*MelM (**Fig 2a**). As previous studies have suggested that *Wolbachia* density can impact *Wolbachia*-based pathogen inhibition, we also examined relative *Wolbachia* density as calculated using previously described methods in single- and double-fed *w*AlbB and *w*MelM mosquitoes^22,23^. *Wolbachia* density was slightly higher in double-fed *w*AlbB mosquitoes, than in single-fed mosquitoes whereas no difference was detected in *Wolbachia* density between single- and double-fed *w*MelM mosquitoes (**Fig 2b**). When individual mosquitoes were examined, there was no correlation between DENV-2 and *Wolbachia* levels for single- and double-fed *w*AlbB mosquitoes (**Fig S1a**). This trend also held true for single- and double-fed *w*MelM mosquitoes (**Fig S1b**). This lack of a strong link between DENV-2 levels and *Wolbachia* density led us to more closely investigate whether the differences in DENV-2 levels between single- and double-fed mosquitoes were due to dissemination status or *Wolbachia* density. Given that we measured DENV-2 infection by testing the mosquito carcass absent the legs and wings, DENV-2 particles that disseminated to other susceptible tissues outside of the midgut would also be included in these measurements. Investigating further, we found that when *w*AlbB mosquitoes were sorted by dissemination status, mosquitoes with a disseminated infection were found to have a higher level of DENV-2 than those with a non-disseminated infection (**Fig 2c**). Additionally, there were no differences in *Wolbachia* densities between *w*AlbB mosquitoes that were not infected, or with either non-disseminated or disseminated infections (**Fig 2d**). Though numbers were limited, we saw similar trends in *w*MelM mosquitoes where mosquitoes with a disseminated infection had higher levels of DENV-2 (**Fig S2a**). As with *w*AlbB mosquitoes there was no difference in *Wolbachia* density between *w*MelM mosquitoes that were either not infected, were infected but had a non-disseminated infection, or had a disseminated infection (**Fig S2b**). Thus, increased dissemination rates were associated with increased DENV-2 levels absent large differences in *Wolbachia* density likely due to earlier escape from the midgut resulting in more opportunities for replication throughout the mosquito body.

**Figure 2:**
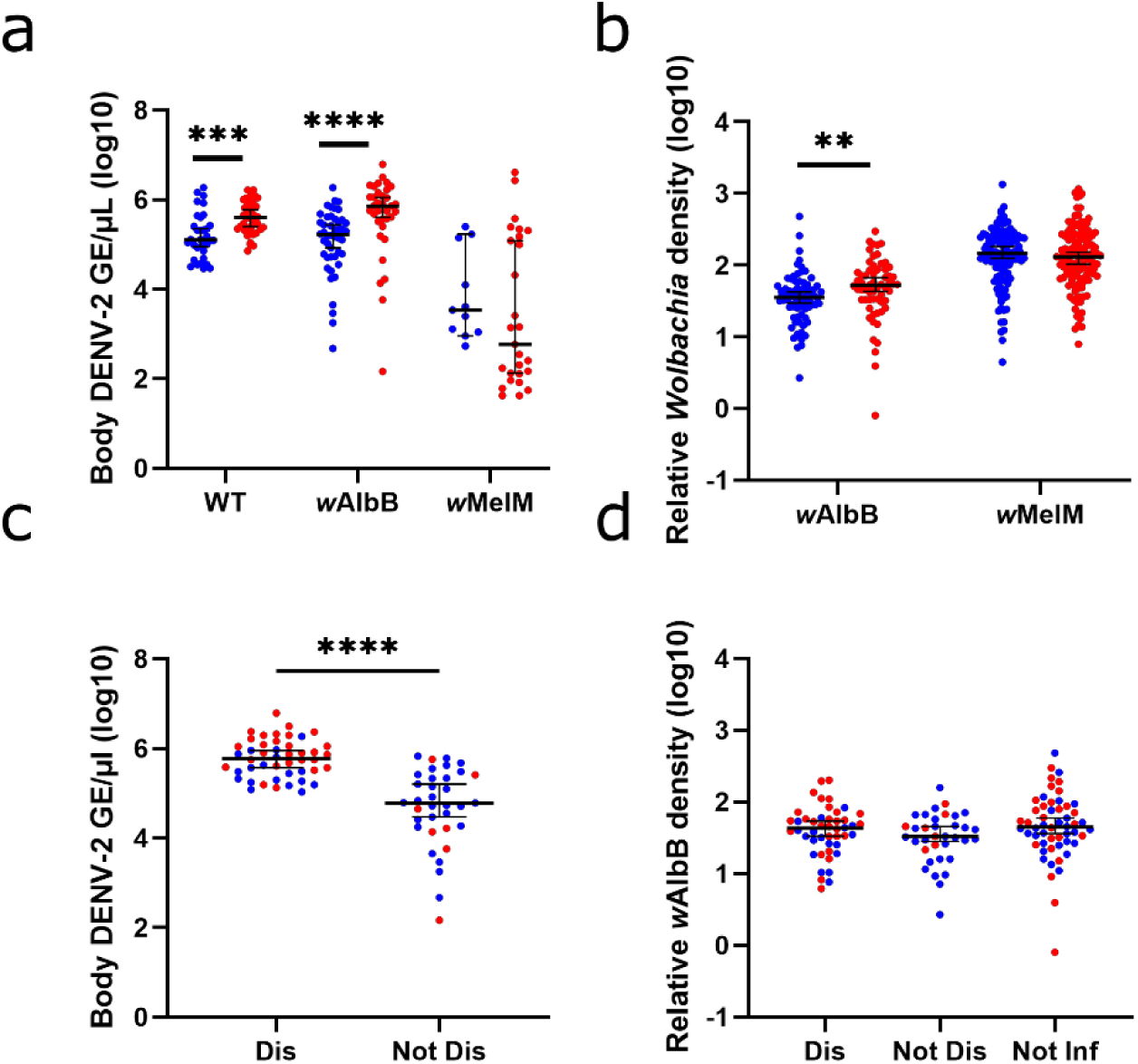
Comparison of DENV-2 titers and relative *Wolbachia* densities at 7 dpi by feeding and dissemination status. **a)** Body DENV-2 titers in single- and double-fed WT, *w*AlbB and *w*MelM mosquitoes. **b)** Relative body *Wolbachia* density in single- and double-fed *w*AlbB and *w*MelM mosquitoes. **c)** Body DENV-2 titers in *w*AlbB mosquitoes by dissemination status and feeding status. **d)** Relative body Wolbachia densities in *w*AlbB mosquitoes by infection, dissemination, and feeding status. Comparisons were made using Mann-Whitney U tests (**a**, **b**, and **c**) or a Kruskal-Wallis test with Dunn’s multiple comparisons (**d**). * = p ≤ 0.05, ** = p ≤ 0.01, *** = p ≤ 0.001, **** = p < 0.0001. Blue = single-fed, red = double-fed. GE = genome equivalents. Dis = mosquitoes with a disseminated infection (i.e., dengue infected bodies and legs), Not Dis = mosquitoes with a non-disseminated infection (i.e., dengue infected bodies, but not legs), Not Inf = mosquitoes that were exposed but are not infected with DENV-2. Lines indicate median with 95% confidence interval.

In our previous studies, we found that multiple blood meals not only increased dissemination rates, but also shortened the EIP^10,16^. Further, we demonstrated that forced salivation assays were less accurate at predicting transmission than using the presence of virus in mosquito legs (dissemination) as a proxy for transmission ability^24^. To determine if a similar temporal shift in dissemination and subsequent EIP occurs in *Ae. aegypti* mosquitoes transinfected with *w*AlbB *Wolbachia*, we conducted time course assays examining infection and dissemination in single- and double-fed WT and *w*AlbB mosquitoes across days 5-10 post infectious blood meal (**Fig 3a**). We did not include *w*MelM mosquitoes in these assays due to the very low rate of infection and dissemination observed in initial experiments despite substantial numbers (**Fig 1b-c**). Our experiments with WT mosquitoes resulted in similar infection rates over time between groups (**Fig 3b**) and significantly earlier dissemination (i.e., shorter EIP) in the double-fed group as compared to the single-fed group, as we previously observed with other virus-vector pairings (**Fig 3c**)^10,13,24^. Importantly, we observed similar patterns for infection and dissemination in mosquitoes with *w*AlbB with similar infection rates between single- and double-fed groups at all timepoints (**Fig 3d**), and a shorter EIP due to increased dissemination during the earlier timepoints (**Fig 3e**). These results are consistent with our previous studies indicating that successive feeding leads to earlier dissemination from the midgut and suggests that the presence of *Wolbachia* does not disrupt this phenotype^10,16^. When we examined *Wolbachia* density over time in single- and double-fed mosquitoes, we did not observe any clear differences in relative *w*AlbB densities in bodies (**Fig S3a**) or midguts (**Fig S3b**) at any of the timepoints following blood feeding. In conclusion, successive blood feeding results in a shorter EIP due to increased dissemination at earlier timepoints in both the presence and absence of *Wolbachia*.

**Figure 3:**
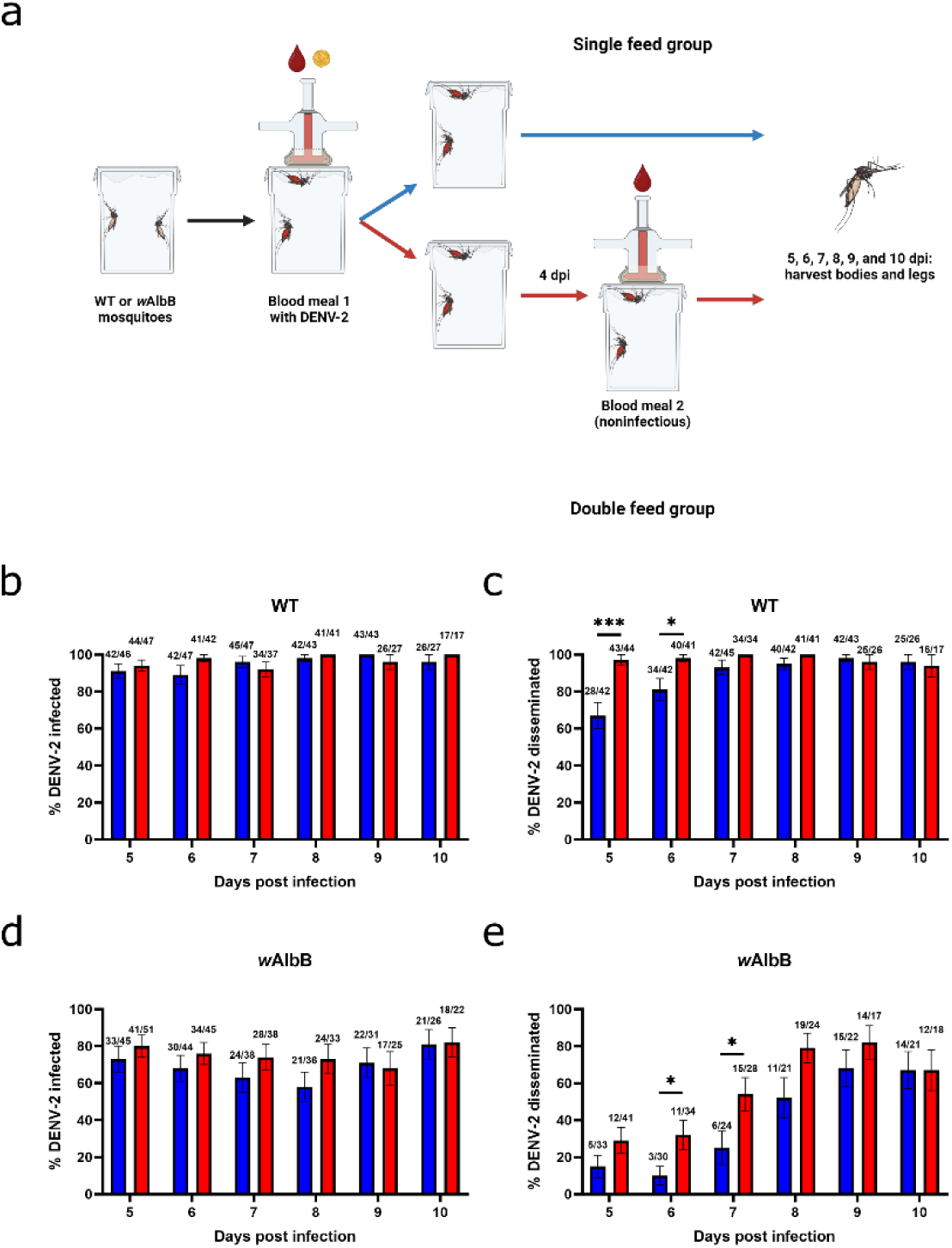
Successive feeding accelerates DENV-2 dissemination and shortens the extrinsic incubation period in both WT and *w*AlbB *Wolbachia* transinfected *Ae. aegypti*. **a)** Experimental design for infection and dissemination time course studies of single- and double-fed WT and *w*AlbB mosquitoes. **b)** Percentage of infected single- and double-fed WT mosquitoes 5-10 dpi. **c)** Percentage of single- and double-fed WT mosquitoes with disseminated infection 5-10 dpi. **d)** Percentage of infected single- and double-fed *w*AlbB mosquitoes 5-10 dpi. **e)** Percentage of single- and double-fed *w*AlbB mosquitoes with disseminated infection 5-10 dpi. Comparisons were made using Fisher’s exact tests. * = p ≤ 0.05, ** = p ≤ 0.01, *** = p ≤ 0.001, **** = p < 0.0001. Blue = single-fed, red = double-fed. Lines indicate mean **±** standard error of the mean.

To accurately estimate the time at which 50% of WT and *w*AlbB mosquitoes have disseminated infections, we used empirical data from our time course experiments to model DENV-2 dissemination (**Fig 4**). As described in the methods section, we used survival models assuming gamma-distributed dissemination times in which the shape (*α*), and rate (β) of the gamma-distribution of DENV-2 dissemination might differ as a function of *w*AlbB infection status (*w*) and blood-feeding status (*f*). As an initial exploration, we fitted four different models: 1) one with four sets of *α*_*w,f*_ and β_w,f_ parameters for each combination of *w* and *f*; 2) one with two sets of *α*_*w*_ and β_w_ parameters for each *w*; 3) one with two sets of *α*_*f*_ and β_f_ parameters for each *f*; and 4) one with a single set of *α* and β parameters. When these models were compared in a pairwise fashion using Bayes factors, the first model that used different dissemination time distributions for all four types of mosquitoes (*w*AlbB/WT x SF/DF) fit the data best, with a Bayes factor of BF_1>2_ = 2.3×10^4^ when compared to the second-best model (**Table S1**). Additional comparisons between models and graphs of prior and posterior shape and rate parameters can be found in supplemental data (**Table S1** and **Fig S4**). From the model of best fit, we calculated the number of days it would take for 50% of mosquitoes to develop a disseminated infection. For single-fed *w*AlbB mosquitoes, 50% dissemination was reached at 8.38 days post-infection (95% credible interval [CrI]: 7.72-9.01; **Fig 4a**). Double-fed *w*AlbB mosquitoes reached 50% dissemination earlier at 6.86 days post-infection (95% credible interval [CrI]: 6.03-7.62; **Fig 4b**). Dissemination in WT mosquitoes was also explored but our modelling was not able to determine a day when 50% of mosquitoes would have a disseminated infection as 50% dissemination was exceeded at every timepoint examined (**Fig 4c** and **4d**).

**Figure 4:**
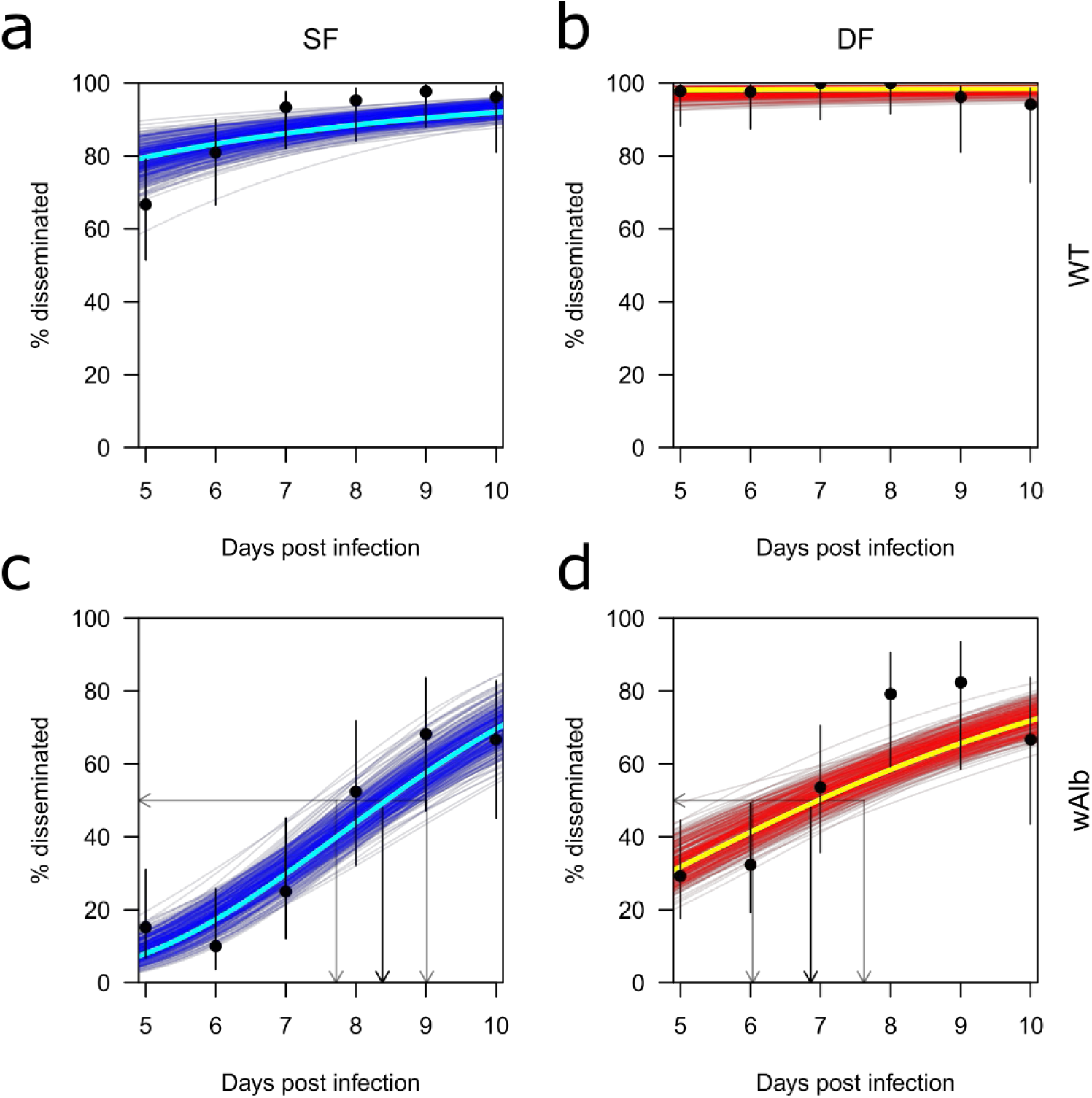
Modeling of timing of dissemination in single and double-fed WT and *w*AlbB *Wolbachia* mosquitoes. Modelling of timing of dissemination for **a)** single-fed WT mosquitoes, **b)** double-fed WT mosquitoes, **c)** single-fed *w*AlbB mosquitoes, and **d)** double-fed *w*AlbB mosquitoes. Experimental data are shown as black dots, with black lines indicating the raw uncertainty in the proportions disseminated. Dark blue or red lines show different draws from the posterior distribution of parameters and indicate the model’s uncertainty. Bright blue and red lines represent the model’s maximum a posteriori prediction. Black lines with arrows mark the timing of 50% dissemination with 95% credible intervals marked by flanking gray lines with arrows. Blue = single-fed, red = double-fed. SF = single-fed, DF = double-fed.

Using our model predictions of dissemination time as an estimate of EIP, we went on to predict the probability of mosquitoes surviving the EIP given varying average lengths of mosquito lifespan (**Fig 5a**). In all instances, double-fed WT or *w*AlbB mosquitoes were more likely to survive the EIP than single- fed counterparts (**Fig 5a**). WT mosquitoes were more likely to survive the EIP than *w*AlbB mosquitoes, regardless of number of blood meals, indicating that *w*AlbB inhibits DENV-2 infection and dissemination (**Fig 5a**). In general, the probability of mosquitoes surviving the EIP increased as average mosquito lifespan increased; however, WT double-fed mosquitoes were highly likely to survive the EIP regardless of average mosquito lifespan (**Fig 5a**). We went on to quantify the epidemiological significance of these factors by calculating the associated odds ratio of surviving the EIP, associated with *w*AlbB infection status and how that effect was modulated by blood-feeding status (**Fig 5b**). For both single- and double-fed mosquitoes, the odds-ratio of surviving EIP increased as average mosquito lifespan increased, indicating that mosquitoes with *w*AlbB were less likely to survive to transmit DENV-2 when lifespan was shorter (**Fig 5b**). This reflects the increase in time to dissemination observed in *w*AlbB mosquitoes relative to WT mosquitoes. The odds-ratios for double-fed mosquitoes were much smaller than those of single-fed mosquitoes, indicating that although successive feeding did reduce EIP for *w*AlbB mosquitoes, successive feeding has a larger impact on EIP in WT mosquitoes (**Fig 5b**). Overall, this suggests that estimates of DENV-2 inhibition by *Wolbachia* based on traditional single-feeding experiments that fail to account for *Ae. aegypti* successive feeding behavior may underestimate the effectiveness of *Wolbachia*.

**Figure 5:**
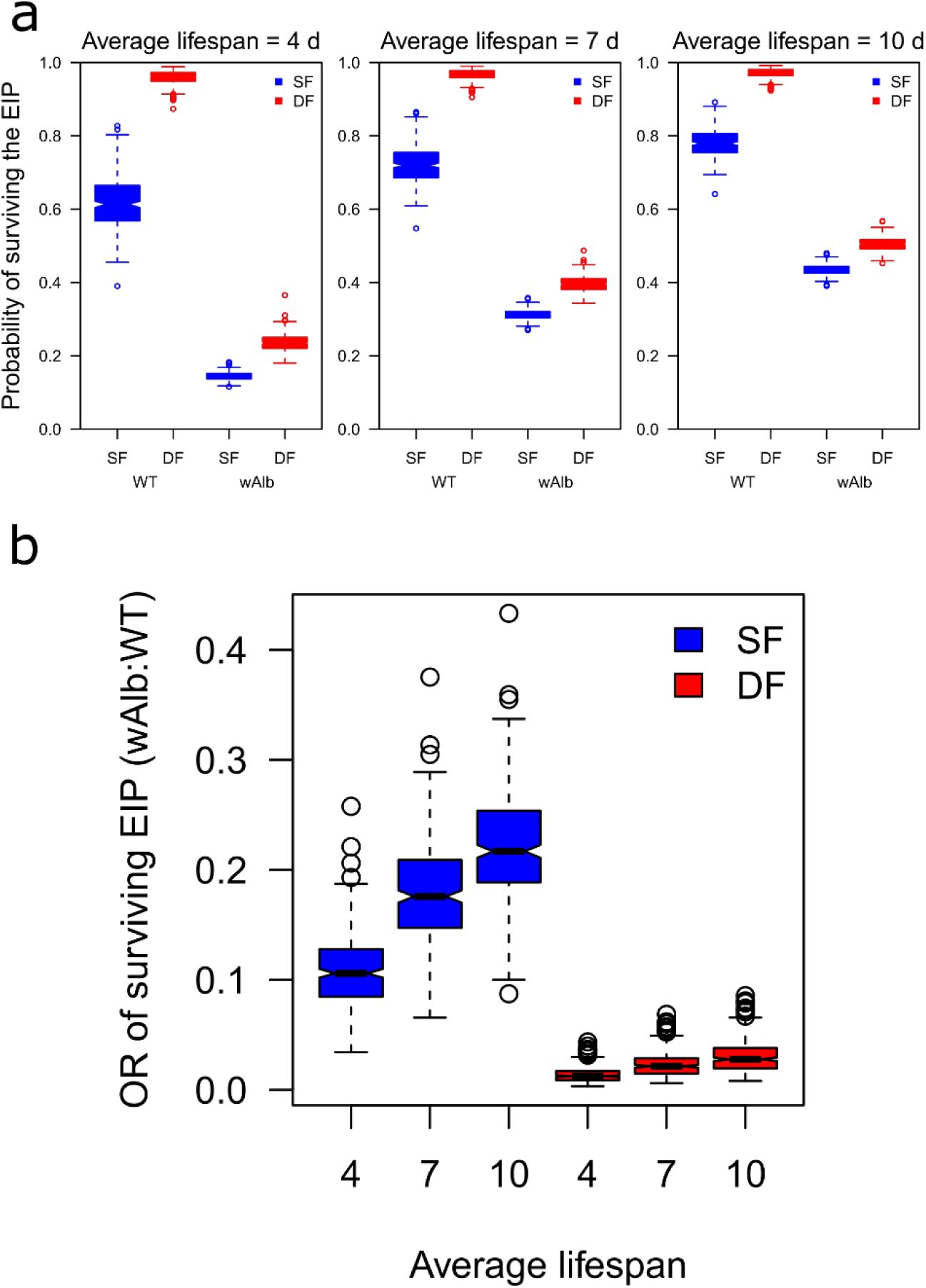
Impact of feeding behavior on the probability of a mosquito surviving the extrinsic incubation period (EIP). **a)** Probability of single- and double-fed WT and *w*AlbB mosquitoes surviving the EIP assuming an average mosquito lifespan of 4, 7, or 10 days **b)** Odds ratio of surviving the EIP (*w*AlbB:WT) given a mosquito lifespan of 4, 7, or 10 days. Boxes and whiskers indicate 50% and 95% posterior prediction intervals, respectively. Circles indicate posterior samples falling beyond the 95% posterior prediction intervals. Blue = single-fed, red = double-fed. SF = single-fed, DF = double-fed.

## Discussion

This work provides important new insights into how mosquito blood feeding behavior impacts the efficiency of *Wolbachia*-based DENV-2 inhibition. In this study, we expanded on our prior work and demonstrated increased early DENV-2 dissemination and shorter EIP in both the presence and absence of *Wolbachia* when *Ae. aegypti* mosquitoes were given a second non-infectious blood meal. By modeling the odds ratio of surviving the EIP, we show that the impact of successive feeding is larger on WT mosquitoes as compared to *w*AlbB mosquitoes, suggesting that *Wolbachia* remains an effective strategy to inhibit DENV-2 transmission even under successive feeding conditions.

We found that WT mosquitoes were much more susceptible to DENV-2 infection than mosquitoes with either *w*MelM or *w*AlbB, indicating strong virus inhibition with *Wolbachia* as has been shown in previous studies (**Fig 1b**)^3,18,25,26^. Further, fewer mosquitoes with *w*MelM became infected than those with *w*AlbB, indicating that *w*MelM provides stronger DENV-2 inhibition or that more of the inhibitory effects of *w*MelM occur prior to DENV-2 infection of the midgut (**Fig 1b**). For all groups, infection rates were largely unchanged by feeding behavior, however, mosquitoes with *w*MelM that were given a second blood meal had a slight increase in infection rate that may be an artifact of the low number of mosquitoes infected (**Fig 1b**, **3b**, and **3d**). Time course experiments revealed that both WT and *w*AlbB mosquitoes had higher rates of early dissemination when given a second blood meal (**Fig 1c**, **3c**, and **3e**). This is in line with earlier results from WT mosquitoes and indicates that despite viral inhibition by *w*AlbB, earlier DENV-2 dissemination in mosquitoes with *Wolbachia* is facilitated by additional non-infectious blood meals^10,13^.

Interestingly, we found that when WT and *w*AlbB mosquitoes were given a second blood meal, there was an increase in DENV-2 levels in the mosquito body (**Fig 2a**). As we found little variation in *Wolbachia w*AlbB or *w*MelM levels by feeding or infection or dissemination status (**Fig 2b**, **2d**, **S1a-d, S2b**, **S3a**, **S3b**), we propose that the differences in virus titer we observed are not correlated with *Wolbachia* density, as has previously been suggested^27^. When DENV-2 levels in *w*AlbB and *w*MelM mosquitoes were compared by dissemination status (**Fig 2c**, **S2a**), mosquitoes with disseminated infections had higher levels of DENV-2, which may indicate that midgut escape allows for invasion of new tissues and increased virus replication or that mosquitoes with higher DENV-2 levels are more likely to become disseminated, as seen in previous studies^16^.

When dissemination time course data from WT and *w*AlbB mosquitoes was used to predict the time needed for 50% of mosquitoes to reach a disseminated infection, we estimated that the incubation periods was approximately 2 days shorter in double-fed as compared to single-fed *w*AlbB mosquitoes (**Fig 3e**, **4c-d**). Experimentally, differences were seen between dissemination rates in WT single- and double-fed mosquitoes, though, predictions of 50% dissemination timing were not possible in WT groups as dissemination was always higher than 50% across the timeframe examined (**Fig 3c**, **4a-b, Table S2**). This is likely due to the starting DENV-2 titer used in our experiments and represents a limitation of our study but also reflects the strong inhibition of infection and dissemination seen in *Wolbachia* transinfected mosquitoes relative to highly susceptible WT populations (**Table S2**). Predictions of the probability of mosquitoes surviving the EIP further reinforced this dynamic, as WT mosquitoes from both single- and double-fed groups were more likely to survive the EIP than their *w*AlbB counterparts given any plausible mosquito lifespan (**Fig 5a**). Although successive feeding always increased the probability of surviving the EIP, when comparing the odds ratio of *w*AlbB:WT mosquitoes surviving the EIP, it became evident that successive feeding increases the probability of WT mosquitoes surviving the EIP more than it does in mosquitoes with *w*AlbB (**Fig 5b**). Given this, *Wolbachia* may have an even stronger impact than previously thought when the sequential feeding behavior of *Ae. aegypti* in the wild is taken into consideration, however, this model does not consider other indirect impacts of *Wolbachia* on mosquito life history traits such as lifespan, fecundity, and feeding frequency^28–31^.

While these findings represent valuable new information regarding virus dynamics in mosquitoes with *Wolbachia*, our study has some limitations. First, although this study attempts to provide a more accurate model of mosquito behavior and feeding in the wild, laboratory experiments are inherently artificial and, in this case, relied on water-jacketed membrane feeders and defibrinated sheep blood rather than a live host with a functioning immune system. We also provided mosquitoes with complete blood meals whereas mosquitoes in the wild frequently take partial blood meals and the viral titers we used to infect mosquitoes may be higher than often encountered in the wild^15,32^.

Additionally, many of our findings are based on data from *w*AlbB mosquitoes as *w*MelM was highly efficient at disrupting DENV-2 infection and subsequent dissemination and we were not able to detect a difference in dissemination rates between single- and double-fed *w*MelM groups due to low numbers (**Fig 1c**). Despite this, the observed trends in *w*AlbB mosquitoes are important to take into account when modelling or considering *Wolbachia*-based interventions and should be examined further using other *Wolbachia* strains and host genetic backgrounds^33^. One finding that should be addressed is the low percentage of *w*AlbB mosquitoes with detectable DENV-2 in their saliva when examined 10 dpi despite adequate levels of infection and dissemination and elevated DENV-2 levels with double-feeding as observed in experiments conducted at earlier timepoints (**Fig 1b-c, 2a, 3d-e, S5a-e**). These assays used forced salivation techniques and, although detection of virus in mosquito saliva can be used to measure transmission ability, previous work has suggested that such artificial salivation assays often underestimate transmission ability and that dissemination as measured by taking mosquito legs more closely reflects the ability of a mosquito to pass on infection^24^. As such, we used dissemination as an estimate of EIP and trust these findings over the limited results from the forced salivation assays.

Despite these limitations, our work provides important new insights into the impact of mosquito feeding behavior on DENV-2 inhibition by *Wolbachia* that will be valuable for future modelling and control efforts. Most prior studies of *Wolbachia*-mediated virus inhibition have not considered the tendency of *Ae. aegypti* to feed frequently and have instead relied on experiments using a single infectious blood meal. Our results indicate that successive blood feeding can impact DENV-2 dissemination timing and the subsequent probability of both WT and *Wolbachia* transinfected mosquitoes surviving the EIP. While our work found increased dissemination with successive feeding in WT mosquitoes and those with *w*AlbB, we also found robust DENV-2 inhibition by *Wolbachia* in both *w*AlbB and *w*MelM groups, with *w*MelM exhibiting even stronger inhibition than *w*AlbB. Our modelling suggests that functional *w*AlbB inhibition of DENV-2 may be even stronger than previously thought due to the larger impact of successive feeding on EIP survivability in WT mosquitoes when compared to *w*AlbB mosquitoes. These results stress the importance of considering mosquito behavior when designing laboratory experiments or modelling control efforts and provide a clearer understanding of DENV-2 infection dynamics in mosquitoes with *Wolbachia* under single- and successively fed conditions.

## Methods

### Mosquito rearing

*Ae. aegypti* mosquito lines (WT, *w*MelM, and *w*AlbB) were generated from natively uninfected *Ae. aegypti* collected near Cairns, Queensland, Australia that were kept uninfected (WT) or transinfected with either *w*AlbB or *w*MelM via microinjection^34,35^. All three mosquito colonies were maintained in separate environmental chambers at 26°C, 60-70% relative humidity, and 12:12 light-dark cycle. Larvae were hatched from egg papers in 500 mL of water and two drops of Liquifry No. 1 fish food. After hatching, approximately 250 first-instar larvae were transferred to trays with 1 L of water and fed with Tetramin baby fish food. Pupae were collected and transferred to Bugdorm-1 cages for adults to emerge. Adult mosquitoes were maintained on 10% sucrose and blood-fed with defibrinated sheep blood. Adults that were approximately one week old were sorted into cups of 60 female mosquitoes/cup and maintained on 10% sucrose-soaked pads prior to and following infection with DENV-2.

### Dengue virus

DENV-2 (125270/VENE93; GenBank: PQ852084) was grown in *Ae. albopictus* C6/36 cells in T75 flasks and split at a 1:15 dilution in 10% FBS MEM media. For DENV-2 infections, when cells were 60-80% confluent, growth media was removed and a thawed 250 μL aliquot of DENV-2 was added to flasks along with 3 mL of 10% FBS MEM media. Flasks were placed on a rocking platform for 1 hour before 12 mL of 10% FBS MEM media was added for a total volume of 15 mL. Infected cells were grown in a 28°C incubator with 5% CO_2_ for five days. DENV-2 was harvested by removing the supernatant from cells 5 days post-infection (dpi). Dilutions of virus-containing cellular supernatant in defibrinated sheep blood were fed to mosquitoes during the primary infectious bloodmeal (**Table S2**).

### Mosquito infections, blood feeding, and stock virus quantification

One day prior to the infectious blood meal, sucrose-soaked pads were replaced with water-soaked pads to stimulate blood feeding. Mosquitoes were fed with 1:5 or 1:12 dilutions of DENV-2 and defibrinated sheep’s blood (**Table S2**). Virus titers fed to mosquitoes were quantified by both RT-qPCR and focus forming assay as described previously (**Table S2**)^12^. Mosquitoes were knocked down on ice and blood fed mosquitoes were sorted into containers and provided with an oviposition cup. Mosquitoes in the double-feed group were given a second, noninfectious blood meal of defibrinated sheep’s blood 4 days after the initial infectious blood meal (**Fig 1a**, 3a**, S3a-b, S5a**). Blood-fed mosquitoes were sorted into new containers and given an oviposition cup. Mosquitoes in both the “single-feed (SF)” and “double-feed (DF)” groups were sacrificed at different days ranging from 5-10 days post infectious blood meal.

### Mosquito tissue collections and extractions

For initial experiments assessing infection and dissemination differences between SF and DF groups at 7 dpi (Infection and dissemination rep 1-5), mosquito bodies were separated from legs and wings (**Table S2**). To assess infection, each mosquito body was placed in a separate 2 mL tube with 200 μL of mosquito diluent (1X phosphate buffered saline with 20% heat-inactivated fetal bovine serum, 50 μg/ml penicillin/streptomycin, 50 μg/ml gentamycin, and 2.5 μg/ml amphotericin B) and a copper bead. To assess dissemination, legs and wings were pooled from individual mosquitoes in a 2 mL tube containing 200 μL of mosquito diluent and a copper bead.

For time course experiments comparing the shift in timing of dissemination (Time course rep 1-1), mosquito bodies and legs were harvested at 5-10 dpi (**Table S2**). As before, tissues were collected into tubes containing 200 μL of mosquito diluent and a copper bead.

To examine salivary transmission in *w*AlbB transinfected mosquitoes (Salivation experiments wAlbB), salivation assays were conducted at 10 dpi (**Table S2**). Wings and legs were pooled in tubes containing mosquito diluent as before and saliva was collected by placing the proboscis of each incapacitated mosquito into a 20 μL pipette tip with 5 μL of a 50:50 mix of 50% sucrose and FBS. Mosquitoes were allowed to salivate for 1 hour before bodies were harvested as before and the saliva-containing solution was expelled into a 2 mL tube containing 100 μL mosquito diluent and a copper bead.

All samples were stored at −80°C until homogenization using a Retsch Mixer Mill 400 for 4 minutes at 30 Hz, followed by centrifugation for 5 min at 7000 rcf. Nucleic acid was extracted from homogenate (75 μL) using the ThermoFisher MagMAX viral/pathogen nucleic acid isolation kit and eluted into 75 μL using the KingFisher Flex system.

## DENV RT-qPCR and FFA

Mosquito samples were screened for DENV-2 RNA using the NEB Luna Universal Probe One-Step RT-qPCR Kit on the Bio-Rad CFX-96 touch real-time PCR detection system using previously developed primers (**Table S3**)^36^. PCR conditions were as follows: 55°C for 10 minutes, 95°C for 1 minute, and 40 cycles of 95°C for 10 seconds followed by 55°C for 30 seconds and a plate read. All plates were run with at least one negative extraction control, one negative template control, and a serial dilution of synthetic RNA transcript. All samples that were quantified were run in duplicate on the same plate. Positivity was determined by Ct value; samples with a Ct value below 37 were considered positive.

A subset of mosquito bodies from WT and *w*AlbB mosquitoes (7 dpi) were used to compare DENV-2 genome equivalents per mL as determined via RT-qPCR to viral titers via focus forming assay (FFA) (**Fig S6a**). For FFAs, *Ae. albopictus* C6/36 cells were seeded into 96-well plates at a density of 3×10^5^ cells/well, incubated overnight at 28°C with 5% CO_2_, and infected the following day with 30 μl per well of virus-containing serially diluted mosquito sample for 1 hour at 28°C with 5% CO_2_. Virus-containing supernatant was removed, cells were covered with 100 μl of 1% methylcellulose in in 10% FBS MEM media, and cells were incubated for 3 days at 28°C with 5% CO_2_. Cells were then fixed for 15 minutes at room temperature with 100 μl of 4% formaldehyde in PBS, washed 3 times with 100 μl PBS, permeabilized with 0.2% Triton-X in PBS for 10 minutes at room temperature, washed again 3 times, and 30 μl of mouse anti-flavivirus group antigen antibody from NovusBio D1-4G2-4-15 (4G2) diluted 1:500 in PBS was added to each well. Plates were incubated overnight at 4°C. Plates were then washed 3 times with PBS and then incubated overnight at 4°C with 30 μl of Invitrogen goat anti-mouse IgG (H+L) cross-adsorbed secondary antibody, Alexa Fluor 488 diluted 1:200 in PBS. The following day, plates were washed to remove excess secondary antibody, and foci were counted using a Zeiss Axio Vert.A1 inverted microscope with a 2.5X objective and a FITC filter. As expected, DENV-2 genome equivalents/mL via RT-qPCR for body samples were higher than focus forming units/mL via FFA, yet DENV-2 concentrations via each method were internally consistent and concentrations are correlated between both methods (**Fig S6a**). This provided justification for using RT-qPCR rather than FFA to measure DENV-2 concentration in experiments.

### Wolbachia and Aedes qPCR

Samples were screened for *Wolbachia* genome equivalents and compared to *Ae. aegypti* S6 DNA copies using the NEB Luna Universal qPCR kit on the CFX Connect Real-Time PCR Detection System and protocols and primers were modified from Lau et. al and Lee et. al (**Table S3**)^37,38^. PCR conditions were as follows: 95**°**C for 3 minutes, 40 cycles of 95**°**C for 10 seconds and 60**°**C for 30 seconds with a plate read at the end of each cycle. A melt curve was run from 65**°**C - 95**°**C at a rate of 0.5**°**C every 5 seconds with a plate read every 5 seconds. All plates were run with at least one negative extraction control, one negative template control, and one each of confirmed positive RNA extracts from WT and *w*MelM and *w*AlbB-infected colonies. All samples that were quantified were run in duplicate on the same plate. Positivity was determined by the peak of the melting curve. For the wMwA *Wolbachia* detection assay, WT mosquitoes produced no melt peak, *w*MelM presence resulted in a melt peak at 80.5**°**C, and *w*AlbB produced a melt peak at 78 - 78.5**°**C. *Ae. aegypti* DNA detection served as an *Ae. aegypti* genome copies control and all mosquito groups produced similar melt peaks at 81.5-82.5**°**C. Relative *Wolbachia* densities were determined by taking the average crossing point (Cp) of the *Wolbachia*-specific marker and the average Cp value of the *Ae. aegypti*-specific marker across 2 duplicate wells. The average *Wolbachia*-specific marker Cp was then subtracted from the average *Ae. aegypti*-specific marker Cp and transformed by 2^n^ as described before^23^.

### Experimental data analysis

Comparisons of proportions of DENV-2 infection, dissemination, and saliva positivity were made using contingency analyses with Fisher’s exact tests. DENV-2 and *Wolbachia* concentration differences between single- and double-fed groups were compared on untransformed data using Mann Whitney U tests. Non-parametric Spearman correlation tests were used to test for correlation between DENV-2 and *Wolbachia* concentrations in individual mosquitoes. For comparisons between uninfected mosquitoes, mosquitoes with disseminated infections, and mosquitoes with non-disseminated infections, a Kruskal-Wallis test with Dunn’s post hoc test for multiple comparisons was used on untransformed data. Specific statistic tests are noted in the legend for each graph.

### Analysis and modelling of results

To analyze the effect of feeding status and *Wolbachia* infection on time to dissemination, we performed a survival analysis assuming a gamma-distributed time to dissemination. This allowed the hazard rate to increase over time, consistent with the expectation that dissemination would be unlikely until after some minimum amount of time. More specifically, we modelled the dpi for DENV-2 infection to disseminate to the salivary glands, *D*, as a gamma-distributed random variable, which is defined by a shape parameter, *α*, and a rate parameter, β. Because *D* is a continuous random variable and dissemination status was recorded at a daily resolution, the probability that *D*=*d* is *F*(*d*;*α*,β) for *d*=5 and *F*(*d*;*α*,β)-*F*(*d*-1;*α*,β) for *d*∈{6,7,8,9,10}, where *F*(*d*;*α*,β) is the gamma cumulative distribution function. Out of *N_d_* mosquitoes tested for dissemination on day *d*, *X_d_* were positive. Together, this means that Pr(*X_d_*=*x*)=Binomial(*x*;*N_d_*,Pr(*D*=*d*)), where Binomial refers to the probability mass function of a binomial random variable.

Batches of mosquitoes tested for DENV-2 dissemination were distinguished by their *w*AlbB infection status, *w*, and their blood-feeding status, *f*. We considered the possibility that *α* and β might differ as a function of *w* and *f*. To assess this, we fitted four different models: 1) one with four sets of *α*_*w,f*_ and β*_w,f_* parameters for each combination of *w* and *f*; 2) one with two sets of *α*_*w*_ and β*_w_* parameters for each *w*; 3) one with two sets of *α*_*f*_ and β*_f_* parameters for each *f*; and 4) one with a single set of *α* and β parameters. We compared these models in a pairwise fashion using Bayes factors, which was obtained by taking the ratio of the marginal likelihoods of two models using the marginalLikelihood function in the BayesianTools package in R version 4.3.2^39,40^. A Bayes factor of 10 or greater was taken as evidence of strong support of the model with the higher marginal likelihood^41^.

The likelihood of each model was *L*({*α*_*w,f*_},{β*_w,f_*}|{*X_d,w,f_*})=Π*_d,w,f_* Pr(*X_d_*=*x*). To define priors for the parameters, we referred to two previous studies^10,17^. For *α* and β, we used previously determined joint posterior estimates of *α*_WT,SF_, *α*_WT,DF_, β_WT,SF_, and β_WT,DF_ to define our joint prior distribution for those parameters^10^. Consistent with previous estimates of the effect of *w*MelM on dissemination, we multiplied prior samples of *α*_WT,SF_ and *α*_WT,DF_ by samples from a normal distribution with a mean of 1.276 and a standard deviation of 0.1469 to obtain prior samples of *α*_*w*__AlbB,SF_ and *α*_*w*__AlbB,DF_^17^. Because the estimates from Ye et al., 2015 only describe the effect of *w*MelM on mean time to dissemination and the ratio of the means of two gamma distributions can be described by the ratio of their shape parameters only, we assumed the same prior distributions for β*_w_*_AlbB,SF_ and β*_w_*_AlbB,DF_ as we did for β_WT,SF_ and β_WT,DF_^17^. Due to a lack of prior knowledge about the correlation among shape and rate parameters for WT and *w*AlbB mosquitoes, we scrambled the ordering between these two sets of parameter samples to make their priors independent.

We obtained posterior estimates of the parameters of each model using the Metropolis sampler in the BayesianTools package in R version 4.3.2^39,40^. We used three chains totaling 105 iterations. We assessed convergence of these chains through visual inspection of trace plots and confirmation that parameter-wise and multivariate Gelman-Rubin diagnostics were all indistinguishable from 1 using the gelmanDiagnostics function in the BayesianTools package in R version 4.3.2^39,40^.

To assess the epidemiological significance of our estimates, we used model predictions of dissemination time to calculate model predictions of the probability of a mosquito surviving long enough for virus to disseminate to the legs or Pr(survive to disseminate). The closely related probability of surviving the EIP is a key quantity in the canonical Ross-Macdonald theory of mosquito-borne pathogen transmission and our earlier work has established that dissemination is an accurate predictor of transmission ability and thus EIP^24,42^. We focused on this metric, because EIP is the only parameter in the Ross-Macdonald expression for the basic reproduction number, R_0_, affected by blood-feeding status, and it is also expected to be strongly affected by *w*AlbB infection status^42^. Thus, the relative effect of blood-feeding status and *w*AlbB infection status on Pr(survive EIP) is identical to their relative effect on R_0_. Because dissemination time is a random variable in our analysis, we calculate an expected value of Pr(survive to disseminate) as

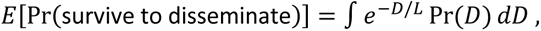

Where *D* is dissemination time, Pr(*D*) is the gamma distribution for *D* estimated above, and *L* is average mosquito lifespan (we explored values of 4, 7, and 10 days). In addition to this qunatity, we also calculated the epidemiological significance of blood-feeding status and *w*AlbB infection status these by calculating the associated odds ratio, OR(survive to disseminate), associated with *w*AlbB infection status and how that effect was modulated by blood-feeding status.

## Data availability

All data are included in this manuscript, the supplementary files, source data, and on GitHub at https://github.com/TAlexPerkins/doubleFeedWolbachia.

## Acknowledgements

We would like to thank Dr. Ary Hoffman for providing resources critical for mosquito colony production. This publication was made possible by CTSA Grant Number UL1 TR001863 from the National Center for Advancing Translational Science (NCATS), a component of the National Institutes of Health (NIH) awarded to CBFV, the National Institute of Allergy and Infectious Diseases of the NIH under award number AI148477 (RMJ & DEB), the NIH T32AI055403 (AS), the National Science Foundation Graduate Research Fellowship under Grant No. DGE-2139841 (AS), NIH National Institute of General Medical Sciences R35 MIRA program to TAP grant number R35GM143029 (AP), Richter Fellowship from Trumbell College (BN), and the Ambrose Monell Foundation (CBFV). PAR was supported by an Australian Research Council Discovery Early Career Researcher Award DE230100067 funded by the Australian Government. The contents of this work are solely the responsibility of the authors and do not necessarily represent the official views of NIH.

## Supplemental data

**Figure S1:**
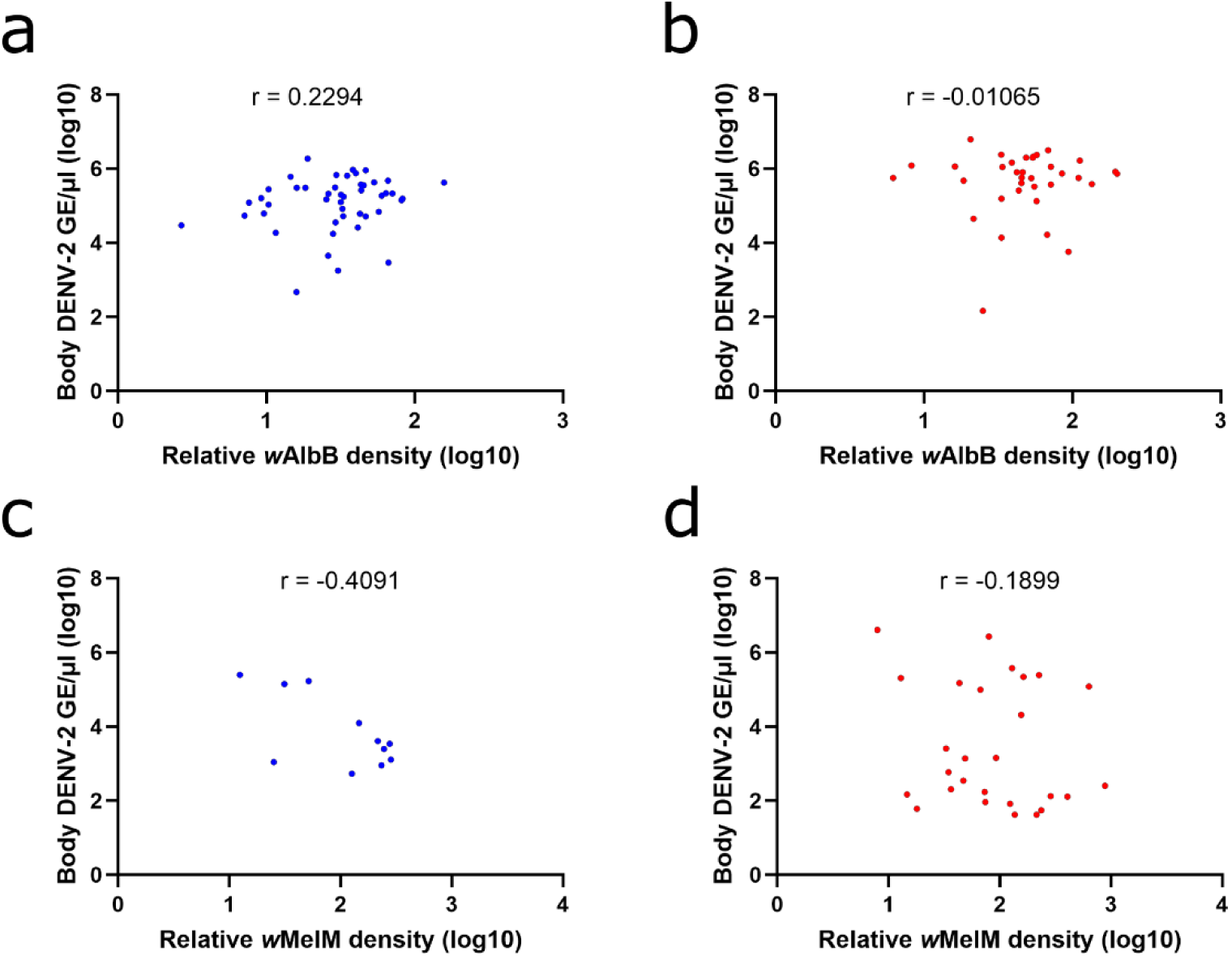
DENV-2 levels vs relative *Wolbachia* densities in bodies from individual single- and double-fed *w*AlbB and *w*MelM mosquitoes. **a)** No correlation for *w*AlbB single-fed mosquito bodies at 7 dpi. **b)** No correlation for *w*AlbB double-fed mosquito bodies at 7 dpi. **c)** No correlation for *w*MelM single-fed mosquito bodies at 7 dpi. **d)** No correlation for *w*MelM double-fed mosquito bodies at 7 dpi. Blue = single-fed, red = double-fed. The correlation between DENV-2 concentration and *Wolbachia* density was compared using non-parametric Spearman correlation for each graph. r = correlation coefficient.

**Figure S2:**
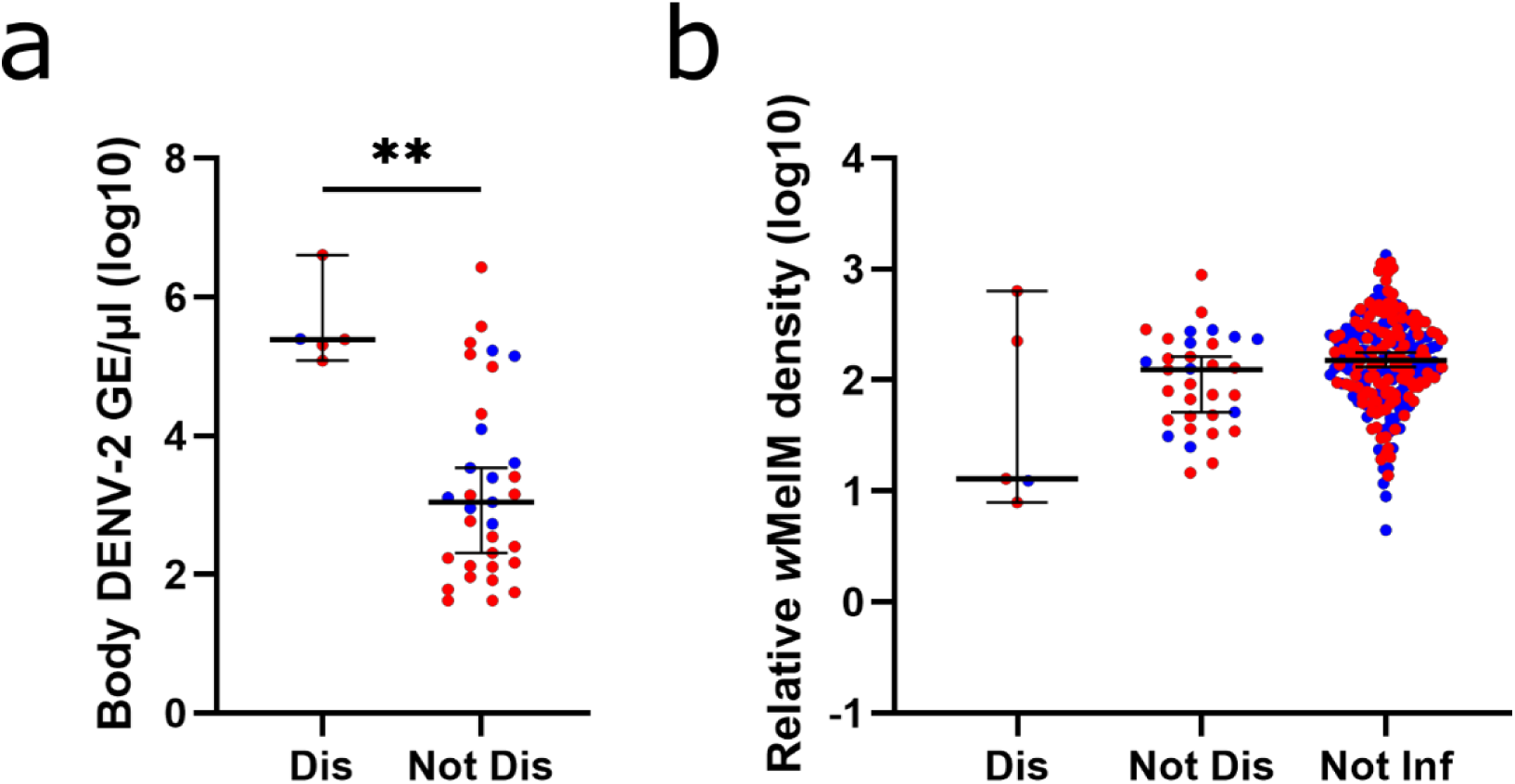
DENV-2 titers and relative *Wolbachia* density in *w*MelM mosquito bodies with disseminated and non-disseminated infection 7 dpi. **a)** Body DENV-2 titers in *w*MelM mosquitoes by feeding status. **b)** Relative body Wolbachia densities in *w*MelM mosquitoes by infection, dissemination, and feeding status. Comparisons were made using a Mann-Whitney U test (**a**) or a Kruskal-Wallis test with Dunn’s multiple comparisons (**b**). * = p ≤ 0.05, ** = p ≤ 0.01, *** = p ≤ 0.001, **** = p < 0.0001. Blue = single-fed, red = double-fed. Lines indicate median with 95% confidence interval.

**Figure S3:**
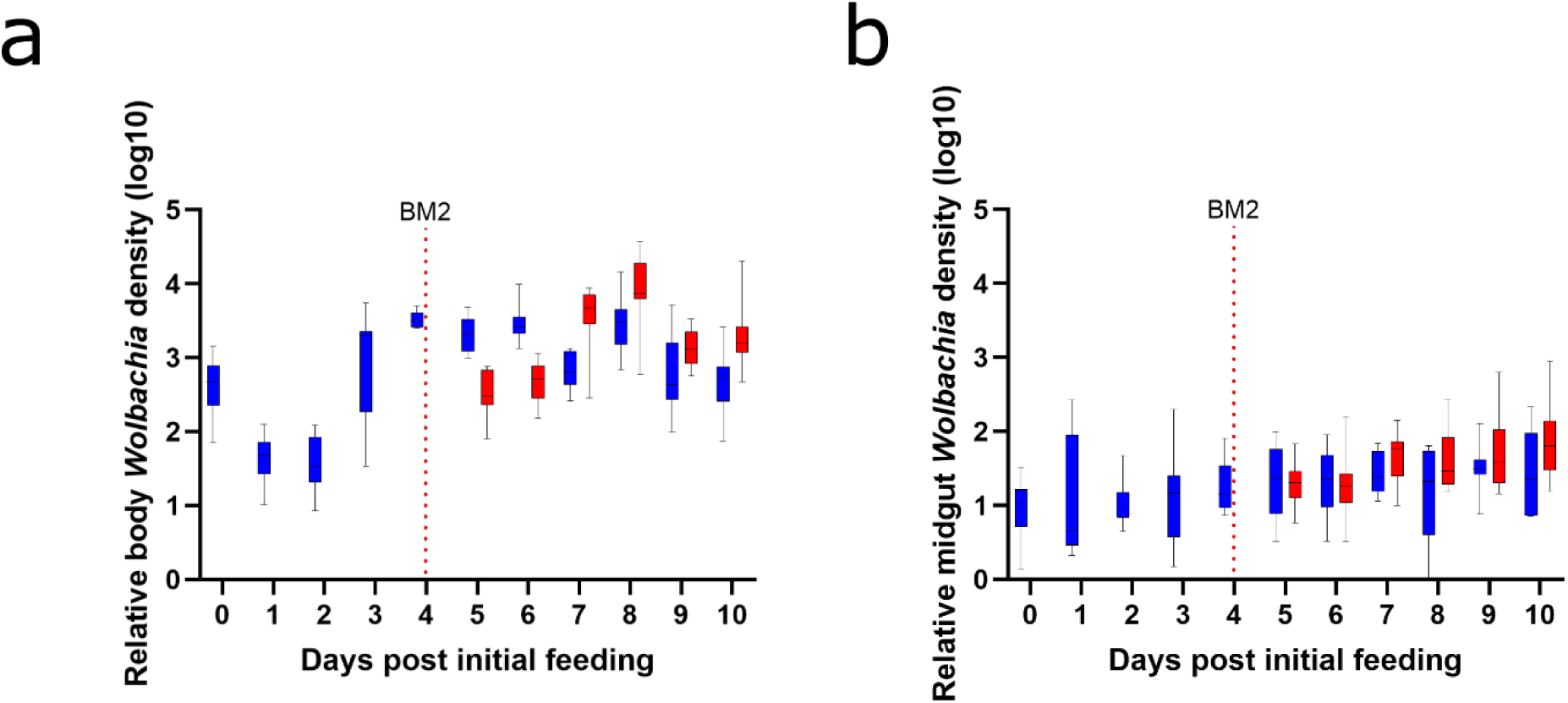
Time course of *Wolbachia* densities in single- and double-fed *w*AlbB mosquitoes. **a)** *w*AlbB Wolbachia densities in single- and double-fed whole mosquitoes following blood feeding. **b)** *w*AlbB Wolbachia densities in single- and double-fed mosquito midguts following blood feeding. Blue = single-fed, red = double-fed. Dashed red lines mark the timing of the second blood meal. Boxes denote first quartile and third quartile with a line at the median. Whiskers indicate minimum and maximum data points.

**Figure S4:**
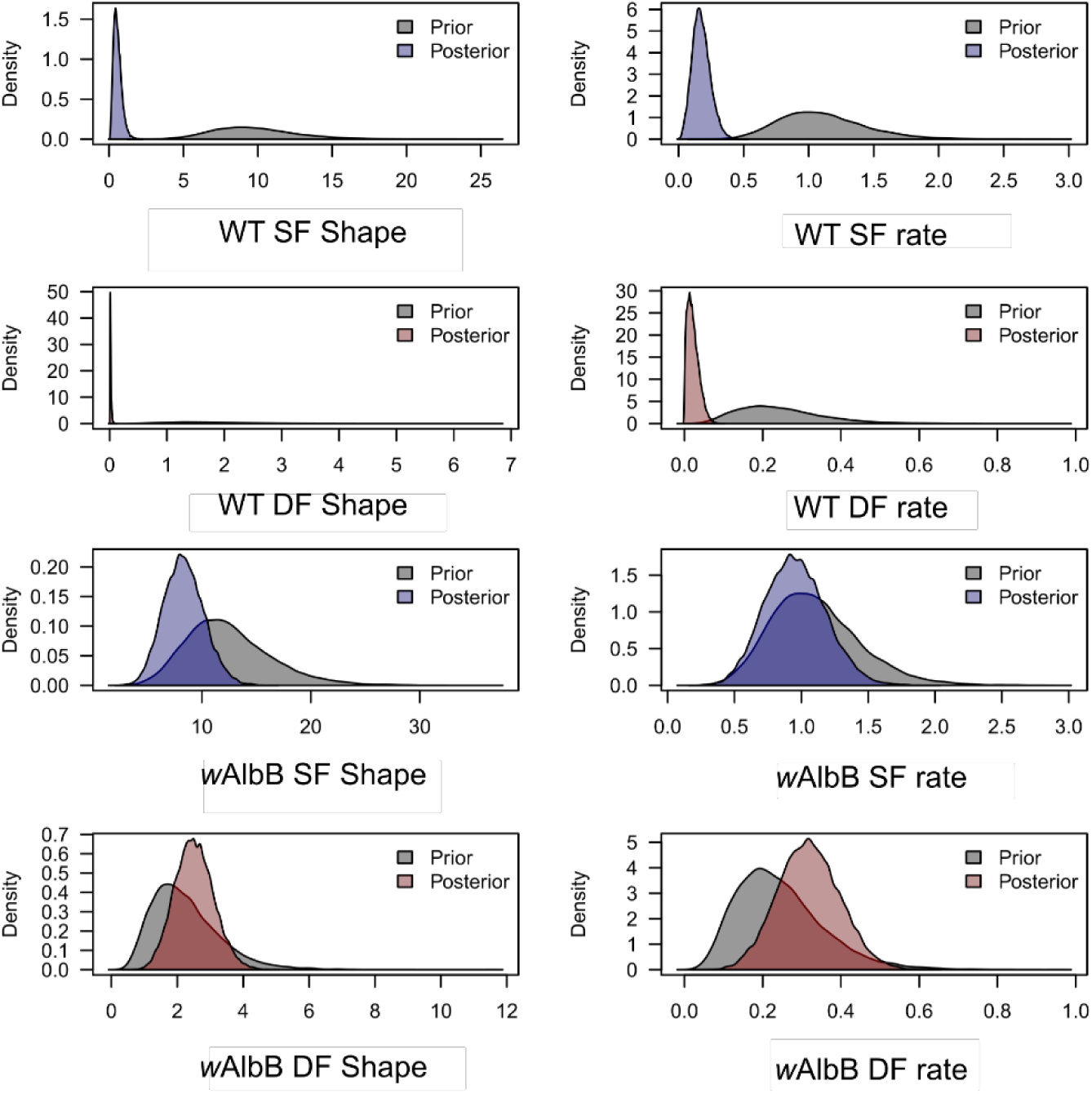
Prior and posterior estimates of shape and rate for gamma-distribution model of dpi for DENV-2 dissemination to the mosquito salivary glands. Blue = single-fed, red = double-fed, gray = prior.

**Figure S5:**
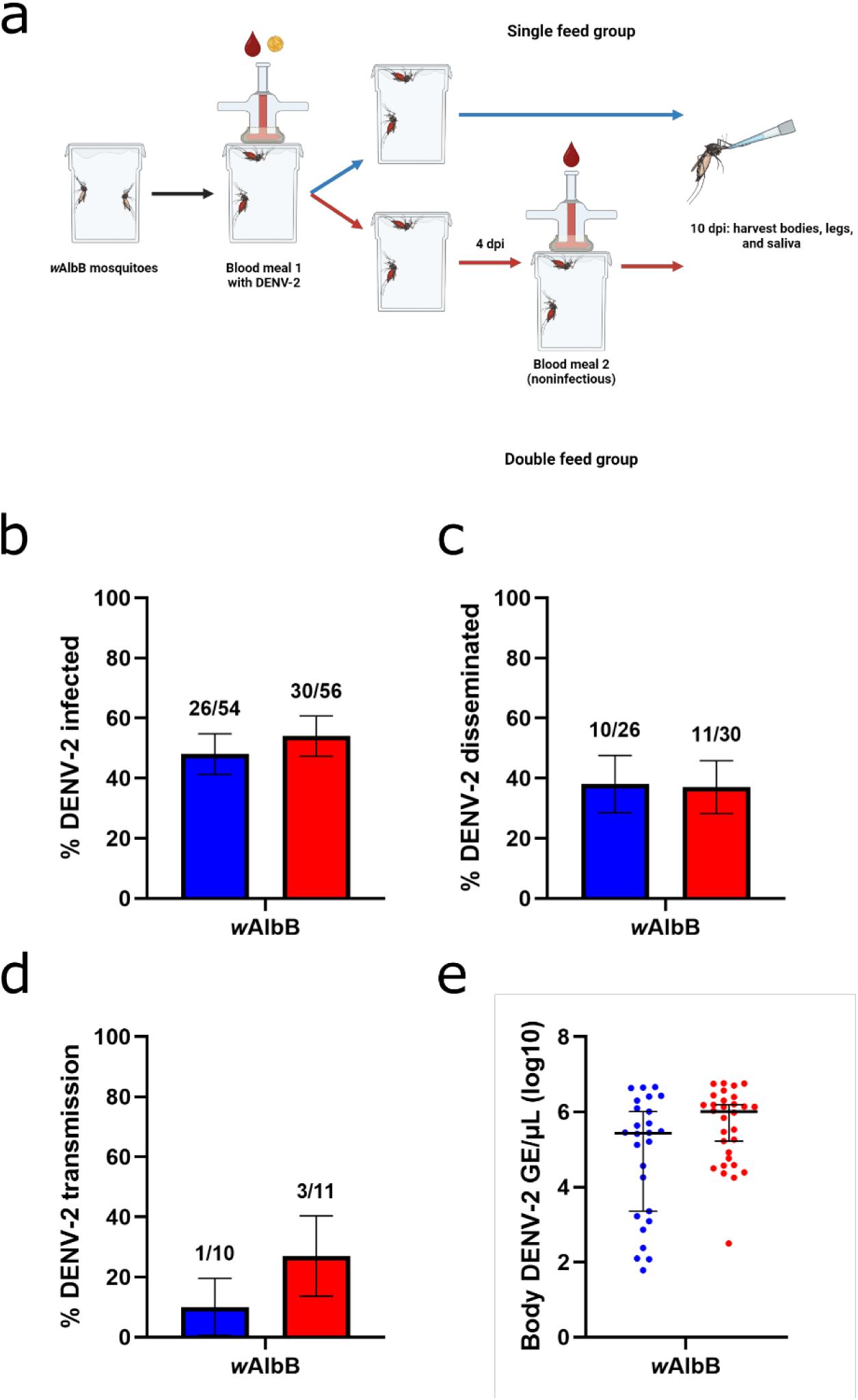
Rates of infection, dissemination, and saliva transmission of DENV-2 in *w*AlbB mosquitoes at 10 dpi. **a)** Experimental design for saliva transmission studies of single- and double-fed *w*AlbB mosquitoes. **b)** Proportion of infected single- and double-fed *w*AlbB mosquitoes at 10 dpi. **c)** Proportion of single- and double-fed *w*AlbB mosquitoes with disseminated DENV-2 infection at 10 dpi. **d)** Proportion of single- and double-fed *w*AlbB mosquitoes with DENV-2 in the saliva at 10 dpi. **e)** DENV-2 titers in single- and double-fed *w*AlbB mosquito bodies at 10 dpi. Comparisons were made using Fisher’s exact tests (**b-d**) or a Mann-Whitney U test (**e**). * = p ≤ 0.05, ** = p ≤ 0.01, *** = p ≤ 0.001, **** = p < 0.0001. Blue = single-fed, red = double-fed. For **b-d**, lines indicate mean **±** standard error of the mean. For **e**, lines indicate median with 95% confidence interval.

**Figure S6:**
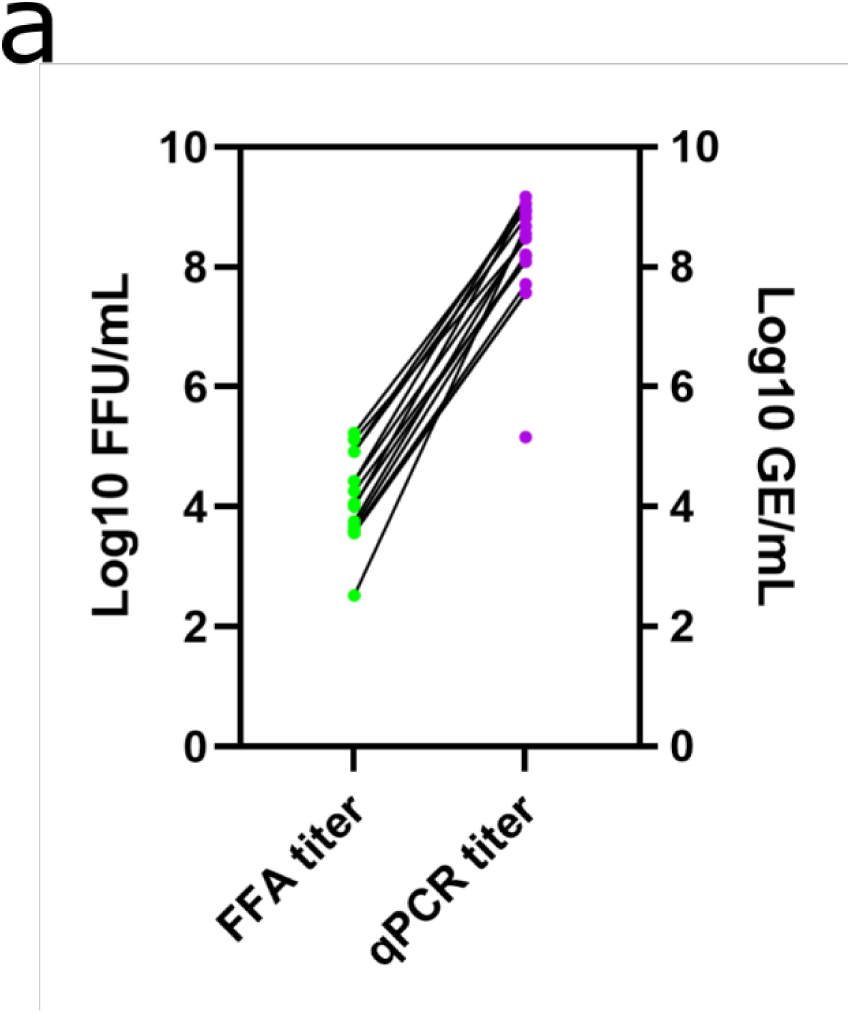
Comparison of DENV-2 concentrations via FFA or RT-qPCR. **a)** Log10 DENV-2 body concentrations in selected samples at 7 dpi as measured by FFA vs measurements through RT-qPCR.

**Table S1:**
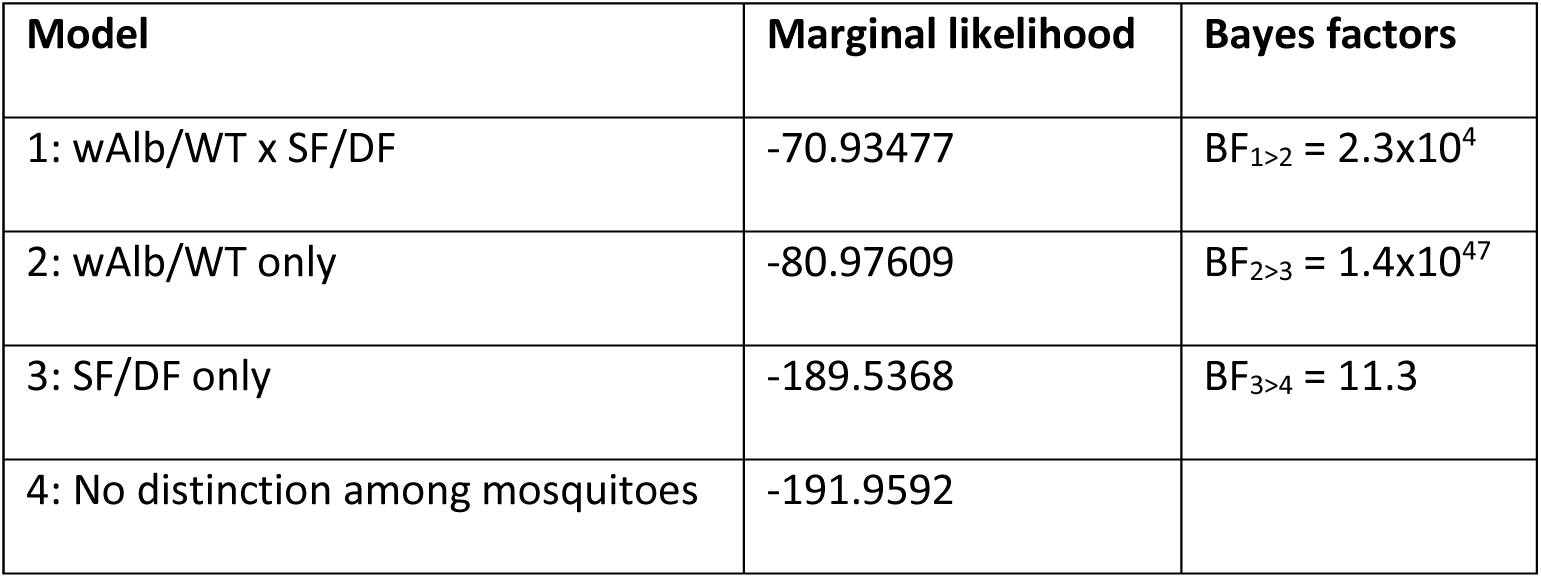
Marginal likelihood and Bayes factors for models of dpi for DENV-2 dissemination to the mosquito salivary glands.

| Model | Marginal likelihood | Bayes factors |
| --- | --- | --- |
| 1: wAlb/WT x SF/DF | -70.93477 | $BF_{1>2} = 2.3 \times 10^4$ |
| 2: wAlb/WT only | -80.97609 | $BF_{2>3} = 1.4 \times 10^{47}$ |
| 3: SF/DF only | -189.5368 | $BF_{3>4} = 11.3$ |
| 4: No distinction among mosquitoes | -191.9592 |  |

**Table S2:** DENV-2 titers given to mosquitoes as measured by qPCR (GE/mL) and FFA (FFU/mL).

| Experiment | Dilution | GE/mL | FFU/mL | Figure showing data |
| --- | --- | --- | --- | --- |
| Infection and dissemination rep 1 | 1:5 | 2.03E+06 | 3.00E+05 | 1, 2, S1, S2, S6 |
| Infection and dissemination rep 2 | 1:5 | 1.67E+06 | 2.00E+05 | 1, 2, S1, S2, S6 |
| Infection and dissemination rep 3 | 1:5 | 2.26E+07 | 1.70E+07 | 1, 2, S1, S2, S6 |
| Infection and dissemination rep 4 | 1:5 | 6.30E+06 | 1.33E+06 | 1, 2, S1, S2, S6 |
| Infection and dissemination rep 5 wMel | 1:5 | 2.28E+07 | 5.00E+06 | 1, 2, S1, S2, S6 |
| Salivation experiments wAlbB | 1:5 | 2.88E+07 | 9.33E+06 | S5 |
| Time course rep 1 | 1:12 | 3.35E+06 | 2.33E+05 | 3, 4 |
| Time course rep 2 | 1:12 | 7.08E+07 | 4.00E+07 | 3, 4 |

**Table S3:**
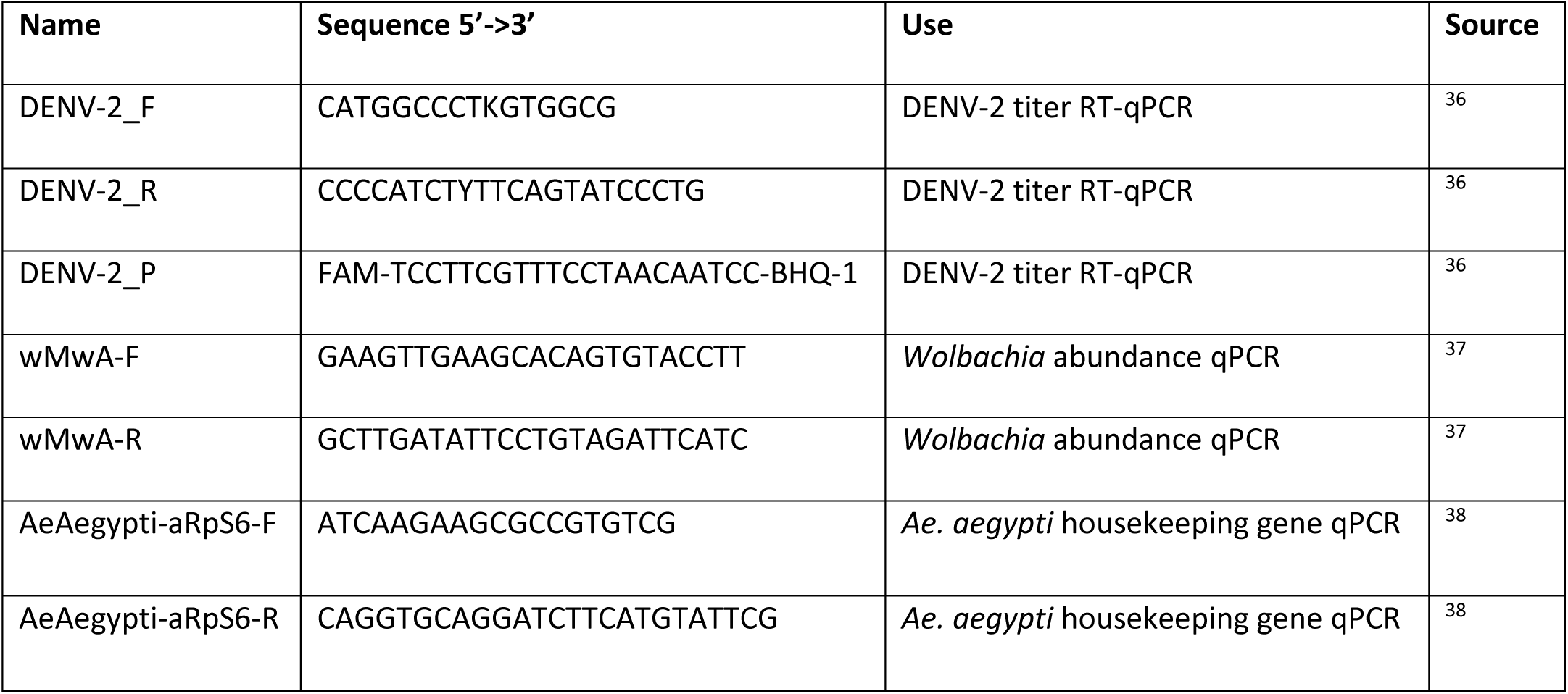
Primers used to measure DENV-2 titers, *Wolbachia* abundance, and *Ae. Aegypti* housekeeping gene copies.

## References

1. 1. World Health Organization. Fact sheets: Dengue and severe dengue. World Health Organization https://www.who.int/health-topics/dengue-and-severe-dengu (2024).

2. World Health Organization. Dengue - Global situation. Disease Outbreak News https://www.who.int/emergencies/disease-outbreak-n https://www.who.int/emergencies/disease-outbreak-news/item/2024-DON518 (2024).

3. Bian, G., Xu, Y., Lu, P., Xie, Y. & Xi, Z. The endosymbiotic bacterium Wolbachia induces resistance to dengue virus in Aedes aegypti. PLoS Pathog. 6, 1–10 (2010).

4. Walker, T. et al. The wMel Wolbachia strain blocks dengue and invades caged Aedes aegypti populations. Nature 476, 450–455 (2011).

5. Ryan, P. A. et al. Establishment of wMel Wolbachia in Aedes aegypti mosquitoes and reduction of local dengue transmission in Cairns and surrounding locations in northern Queensland, Australia. Gates Open Res. 3, 1–32 (2019).

6. Utarini, A. et al. Efficacy of Wolbachia-Infected Mosquito Deployments for the Control of Dengue. N. Engl. J. Med. 384, 2177–2186 (2021).

7. Martinez, J. et al. Genomic and Phenotypic Comparisons Reveal Distinct Variants of Wolbachia Strain wAlbB. Appl. Environ. Microbiol. 88, 1–15 (2022).

8. Crawford, J. E. et al. Efficient production of male Wolbachia-infected Aedes aegypti mosquitoes enables large-scale suppression of wild populations. Nat. Biotechnol. 38, 482–492 (2020).

9. Ant, T. H., Mancini, M. V., McNamara, C. J., Rainey, S. M. & Sinkins, S. P. Wolbachia-Virus interactions and arbovirus control through population replacement in mosquitoes. Pathog. Glob. Health 117, 245–258 (2023).

10. Armstrong, P. M. et al. Successive bloodmeals enhance virus dissemination within mosquitoes and increase transmission potential. Nat. Microbiol. 5, 239–247 (2020).

11. Brackney, D. E., LaReau, J. C. & Smith, R. C. Frequency matters: How successive feeding episodes by blood-feeding insect vectors influences disease transmission. PLoS Pathog. 17, 1–8 (2021).

12. Johnson, R. M., Cozens, D. W., Ferdous, Z., Armstrong, P. M. & Brackney, D. E. Increased blood meal size and feeding frequency compromise Aedes aegypti midgut integrity and enhance dengue virus dissemination. PLoS Negl. Trop. Dis. 17, e0011703 (2023).

13. Ferdous, Z. et al. Multiple bloodmeals enhance dissemination of arboviruses in three medically relevant mosquito genera. Parasit. Vectors 17, 432 (2024).

14. 14. Harrington, L. C., et al. Heterogeneous feeding patterns of the dengue vector, Aedes aegypti, on individual human hosts in rural Thailand. *PLoS Negl. Trop. Dis.* 8, e3048 (2014).

15. Scott, T. W. et al. Detection of multiple blood feeding in Aedes aegypti (Diptera: Culicidae) during a single gonotrophic cycle using a histologic technique. J. Med. Entomol. 30, 94–99 (1993).

16. Johnson, R. M. et al. Investigating the dose-dependency of the midgut escape barrier using a mechanistic model of within-mosquito dengue virus population dynamics. PLoS Pathog. 20, 1–28 (2024).

17. Ye, Y. H. et al. Wolbachia reduces the transmission potential of dengue-infected Aedes aegypti. PLoS Negl. Trop. Dis. 9, 1–19 (2015).

18. Moreira, L. A. et al. A Wolbachia Symbiont in Aedes aegypti Limits Infection with Dengue, Chikungunya, and Plasmodium. Cell 139, 1268–1278 (2009).

19. Amuzu, H. E., Simmons, C. P. & McGraw, E. A. Effect of repeat human blood feeding on Wolbachia density and dengue virus infection in Aedes aegypti. Parasites and Vectors 8, 1–9 (2015).

20. Lau, M. J. et al. The effect of repeat feeding on dengue virus transmission potential in Wolbachia-infected Aedes aegypti following extended egg quiescence. PLoS Negl. Trop. Dis. 18, 1–16 (2024).

21. Ross, P. A. et al. Developing Wolbachia-based disease interventions for an extreme environment. PLoS Pathog. 19, 1–26 (2023).

22. Amuzu, H. E. & McGraw, E. A. Wolbachia-Based Dengue Virus Inhibition Is Not Tissue-Specific in Aedes aegypti. PLoS Negl. Trop. Dis. 10, 1–18 (2016).

23. Ross, P. A. et al. A decade of stability for wMel Wolbachia in natural Aedes aegypti populations. PLoS Pathog. 18, 1–18 (2022).

24. Gloria-Soria, A., Brackney, D. E. & Armstrong, P. M. Saliva collection via capillary method may underestimate arboviral transmission by mosquitoes. Parasites and Vectors 15, 1–9 (2022).

25. Tan, C. H. et al. wMel limits zika and chikungunya virus infection in a Singapore Wolbachia-introgressed Ae. aegypti strain, wMel-Sg. PLoS Negl. Trop. Dis. 11, 1–10 (2017).

26. Caragata, E. P., Dutra, H. L. C. & Moreira, L. A. Inhibition of Zika virus by Wolbachia in Aedes aegypti. Microb. Cell 3, 293–295 (2016).

27. Lu, P., Bian, G., Pan, X. & Xi, Z. Wolbachia induces density-dependent inhibition to dengue virus in mosquito cells. PLoS Negl. Trop. Dis. 6, 1–8 (2012).

28. Lau, M. J., Ross, P. A. & Hoffmann, A. A. Infertility and fecundity loss of wolbachia-infected aedes aegypti hatched from quiescent eggs is expected to alter invasion dynamics. PLoS Negl. Trop. Dis. 15, 1–16 (2021).

29. Lau, M. J., Ross, P. A., Endersby-Harshman, N. M., Yang, Q. & Hoffmann, A. A. Wolbachia inhibits ovarian formation and increases blood feeding rate in female Aedes aegypti. PLoS Negl. Trop. Dis. 16, 1–16 (2022).

30. Maciel-de-freitas, R. et al. Wolbachia strain wMel and wAlbB differentially affect Aedes aegypti Traits Related To Fecundity. Microbiol. Spectr. 12, 1–20 (2024).

31. McMeniman, C. J. et al. Stable introduction of a life-shortening Wolbachia infection into the mosquito Aedes aegypti. Science (80-.). 323, 141–144 (2009).

32. Nguyen, N. M. et al. Host and viral features of human dengue cases shape the population of infected and infectious Aedes aegypti mosquitoes. Proc. Natl. Acad. Sci. U. S. A. 110, 9072–9077 (2013).

33. Liang, X. et al. Wolbachia wAlbB remains stable in Aedes aegypti over 15 years but exhibits genetic background-dependent variation in virus blocking. PNAS Nexus 1, 1–11 (2022).

34. Gu, X. et al. A wMel Wolbachia variant in Aedes aegypti from field-collected Drosophila melanogaster with increased phenotypic stability under heat stress. Environ. Microbiol. 24, 2119– 2135 (2022).

35. Ross, P. A. et al. A wAlbB Wolbachia Transinfection Displays Stable Phenotypic Effects across Divergent Aedes aegypti Mosquito Backgrounds. Appl. Environ. Microbiol. 87, 1–19 (2021).

36. Callahan, J. D. et al. Development and evaluation of serotype- and group-specific fluorogenic reverse transcriptase PCR (TaqMan) assays for dengue virus. J. Clin. Microbiol. 39, 4119–4124 (2001).

37. Lau, M. J., Hoffmann, A. A. & Endersby-Harshman, N. M. A diagnostic primer pair to distinguish between wMel and wAlbB Wolbachia infections. PLoS One 16, 1–11 (2021).

38. Lee, S. F., White, V. L., Weeks, A. R., Hoffmann, A. A. & Endersby, N. M. High-throughput PCR assays to monitor Wolbachia infection in the dengue mosquito (Aedes aegypti) and Drosophila simulans. Appl. Environ. Microbiol. 78, 4740–4743 (2012).

39. Hartig, F., Minunno, F. & Paul, S. BayesianTools: General-Purpose MCMC and SMC Samplers and Tools for Bayesian Statistics. R package version 0.1.8 (2023).

40. R Core Team. R: A Language and Environment for Statistical Computing. R Foundation for Statistical Computing, Vienna, Austria (2023).

41. Jeffereys, H. The Theory of Probability (3rd ed.). (Oxford Unversity Press, 1961).

42. Smith, D. L. et al. Ross, Macdonald, and a theory for the dynamics and control of mosquito-transmitted pathogens. PLoS Pathog. 8, 1–13 (2012).

